# scUnify: a unified framework for training and inference across multiple single-cell foundation models

**DOI:** 10.64898/2026.03.01.708392

**Authors:** Donghee Kim, Ari Hong, Kyeonghun Jeong, Kwangsoo Kim

**Affiliations:** Biomedical Research Institute, Seoul National University Hospital Seoul, Republic of Korea; Interdisciplinary Program in Bioinformatics, Seoul National University, Seoul, Republic of Korea; Interdisciplinary Program in Bioengineering, Seoul National University, Seoul, Republic of Korea; Department of Transdisciplinary Medicine, Seoul National University Hospital, Seoul, Republic of Korea; Department of Medicine, College of Medicine, Seoul National University, Seoul, Republic of Korea; Center for Data Science, Healthcare AI Research Institute, Seoul National University Hospital, Seoul, Republic of Korea

**Keywords:** Single-cell RNA sequencing, Single-cell foundation models, Computational framework, Model adaptation

## Abstract

Single-cell foundation models (scFMs) differ in software requirements and performance across downstream tasks and adaptation strategies, complicating comparison and reuse. We present scUnify, a framework that preserves each backbone’s required processing while separating model-specific trainers, downstream tasks, and adaptation strategies as reusable components. Across five scFMs, scUnify reproduced original inference and training workflows, extended model-native tasks with multiple parameter-efficient fine-tuning methods, and demonstrated extensibility by connecting a newly implemented custom trainable task to multiple backbones and adaptation strategies. Together, these capabilities enable researchers to systematically compare these combinations and extend custom tasks across heterogeneous scFMs within a common workflow.

## 1 Background

The accumulation of large single-cell RNA sequencing (scRNA-seq) atlases has enabled the pretraining of transcriptional patterns observed across tissues and studies and their transfer to new data. Since 2023, various single-cell foundation models (scFMs) that learn transferable cell representations from large atlases have continued to emerge[1]. Early scFMs, including scGPT, Geneformer, and scFoundation, were pre-trained primarily on large scRNA-seq datasets. Those model proofed the applicability to downstream tasks such as celltype classification, batch correction, perturbation prediction, and drug-response prediction[2–4]. Recently, UCE and Nicheformer cover broader biological context into cell representation by exploiting the additional information such as protein-sequence priors embedded by a protein language or spatial transcriptomic expression[5, 6].

However, as the number and scope of scFMs have expanded rapidly, selecting a appropriate model for researcher’s data and research objective has become complex. Moreover, evaluation study of various scFMs across multiple datasets and downstream tasks showed that different scFMs had superior performance depending on the task, and that no single scFM demonstrated consistently dominated performance across all comparisons[7, 8]. It suggests that rating the model suitability is inappropriate from previously reported ranks and that comparing candidate scFMs under the research context is important. However, scFMs differ in model-specific preprocessing and input construction, software environments, and training procedures, and each model is distributed primarily through an independent repository. Consequently, comparing the growing range of scFM options requires a recurring model-specific engineering burden including repeated data preparation and environment setup.

Recent evaluation studies have reported that pretrained cell representations do not consistently outperform simpler methods or task-specific baselines across downstream settings. In particular, performance advantages were inconsistent when frozen cell representations were used directly or with probing[8, 9]. Although Full-FT can improve performance over probing in some settings, supervised fine-tuning evaluations of celltype annotation and perturbation prediction showed that its gains were not consistent and that simpler baselines could remain competitive, while updating all model parameters also incurs substantial computational cost[10, 11]. Meanwhile, scPEFT applied adapter-based parameter-efficient fine-tuning (PEFT) to multiple scFMs and showed that it could achieve performance comparable to or better than probing or Full-FT in some of the evaluated downstream and out-of-distribution settings[12]. These findings establish PEFT as an important adaptation option for scFM applications. However, applying the same PEFT method across scFMs with heterogeneous backbone architectures and training procedures still requires model-specific integration and validation, leaving an implementation burden for systematic adaptation comparisons.

These studies show that researchers need to consider adaptation strategies, including probing, Full-FT, and PEFT, alongside multiple candidate scFMs when evaluating models and training conditions for their downstream tasks. However, because each scFM has its own software environment and downstream task learning procedure and applying PEFT also requires model-specific implementation, the recurring engineering burden grows with the number of combinations among models, downstream tasks, and adaptation strategies considered, creating a need for an integrated framework that can consistently execute and compare multiple models and adaptation conditions across diverse downstream tasks within a common analytical workflow. In response to this need, several scFM integration frameworks have recently been developed to address different aspects of scFM execution, benchmarking, and downstream use (Supplementary Table S1). First, scDrugMap integrates multiple scFMs into a drug-response prediction workflow and a web-based service, allowing users to explore model-specific predictions and benchmark results for drug response[13], whereas scFM-eval provides a containerized workflow for standardized execution and benchmarking of multiple scFMs across diverse datasets under zero-shot, few-shot, and fine-tuning settings and manages multiple runs within the same workflow[14]. Although these tools integrate multiple scFMs within a drug-response application or a general execution and evaluation infrastructure, neither demonstrates support for extending user-defined trainable downstream tasks across multiple backbones or applying and comparing multiple PEFT methods. Moving beyond execution and evaluation, BioLLM connects model loading, cell embedding generation, and selected downstream fine-tuning workflows across multiple scFMs through a common model loader and a downstream task interface termed BioTask, allowing new analytical tasks to be added[15]. Similarly, GB.ModelGenerator connects foundation models spanning the DNA, RNA, protein, and cell domains through a common backbone–task interface and enables custom tasks to be constructed with the GB.Cell family, scFoundation, and Geneformer in the single-cell domain[16]. These two frameworks therefore provide interfaces for extending trainable downstream tasks. However, neither framework demonstrates the common application and comparison of diverse PEFT methods across multiple scFMs. Collectively, existing tools have expanded their scope from scFM execution and benchmarking to extensible trainable downstream tasks, but a single-cell workflow that reuses the same trainable task across heterogeneous scFM backbones while independently applying diverse PEFT methods to compare model–task–adaptation combinations has not yet been fully addressed.

To address this gap, we developed scUnify, a framework that enables researchers to compare multiple scFMs and adaptation strategies for their downstream tasks and extend new trainable tasks across diverse scFMs. Its core design preserves the software environment required by each scFM and encapsulates model-specific preprocessing, input construction, and forward procedures within a model-specific trainer, while treating downstream task learning and adaptation strategies as reusable components separated from the backbone. This separation allows researchers to retain the same task-learning logic while changing the scFM backbone and independently applying and comparing probing, Full-FT, and multiple PEFT methods. Moreover, when adding a new trainable task, researchers need only implement its task-specific learning logic through the common task interface, after which the task can be connected to multiple scFM backbones and adaptation strategies. Beyond supporting these model–task–adaptation combinations, scUnify provides a core system for scheduling and managing the growing number of experiments. When multiple experiments are submitted together, the core system manages run allocation, queuing, execution, and result collection according to the available computational resources, while researchers can monitor overall progress through the runtime dashboard. Each run is executed using an AnnData object and a single configuration file specifying the combination and execution conditions, with cell embeddings and task-specific predictions returned in AnnData format for direct compatibility with the scverse ecosystem widely used in scRNA-seq analysis[17].

In addition, scUnify provides several supporting features that reflect practical workflows for using scFMs in scRNA-seq studies. First, researchers who wish to explore pretrained cell representations before downstream training or use them without additional learning can employ a lightweight inference mode that generates frozen cell embeddings without parameter updates. Second, because cell representations are major outputs of scFMs and their quantitative evaluation is important for subsequent applications, the scUnify Evaluator integrates cell-embedding evaluation metrics from scIB[18] and scGraph[19], allowing representations to be assessed within the framework. Finally, recognizing that preparing the configuration file defining each experiment can itself create a barrier to framework use, scUnify provides assistant skills that direct external LLM-based coding assistants to its public documentation when preparing configurations. In this way, scUnify supports an scRNA-seq research workflow that extends beyond model execution, from configuration preparation to cell-representation generation and evaluation.

scUnify currently supports five scFMs—Geneformer, scGPT, scFoundation, UCE, and Nicheformer—and we evaluated its functionality across six datasets through resource profiling, four built-in downstream tasks comprising celltype classification, regression, batch correction, and perturbation prediction, and a custom MIL task. Resource profiling showed that the scUnify core system maintained consistency with the original workflows in the evaluated numerical outputs and training behavior, while the end-to-end runtime did not increase under the tested conditions. Using built-in tasks that retained the original downstream designs of the supported models, we further demonstrated that multiple PEFT methods could be applied and compared within a common workflow. The custom MIL experiment extended this evaluation to a complex new task, showing that its task-specific learning logic could be implemented once and subsequently connected to five scFM backbones and multiple adaptation strategies. Consistent with previous benchmarking and PEFT evaluation studies[7, 8, 12], no single model–adaptation combination consistently dominated across all evaluated downstream tasks. Together, these findings underscore the need to compare candidate combinations within the researcher’s target context and demonstrate that scUnify provides an effective framework for conducting and extending such comparisons.

## 2 Results

### 2.1 Overview of the scUnify framework

scUnify is a framework that connects model preparation and data processing for heterogeneous single-cell foundation models (scFMs) with inference or downstream training, output storage, and subsequent analysis in a united workflow (Figure 1a). During installation, a separate software environment is automatically prepared for each supported model. Once installed, researchers can run a selected scFM by providing a common AnnData input and a configuration file that specifies the model, execution mode, and analysis conditions. During execution, scUnify selects the corresponding model environment, applies model-specific preprocessing and input construction, and then performs inference or training with optimized computational resources management. The resulting cell embeddings and task-specific outputs are stored in consistent AnnData fields for subsequent analysis in existing Scanpy and scverse workflows. This workflow reduces the need to install and switch model-specific environments manually or repeatedly convert input and output formats when analyzing one dataset with multiple scFMs.

**Fig. 1.**
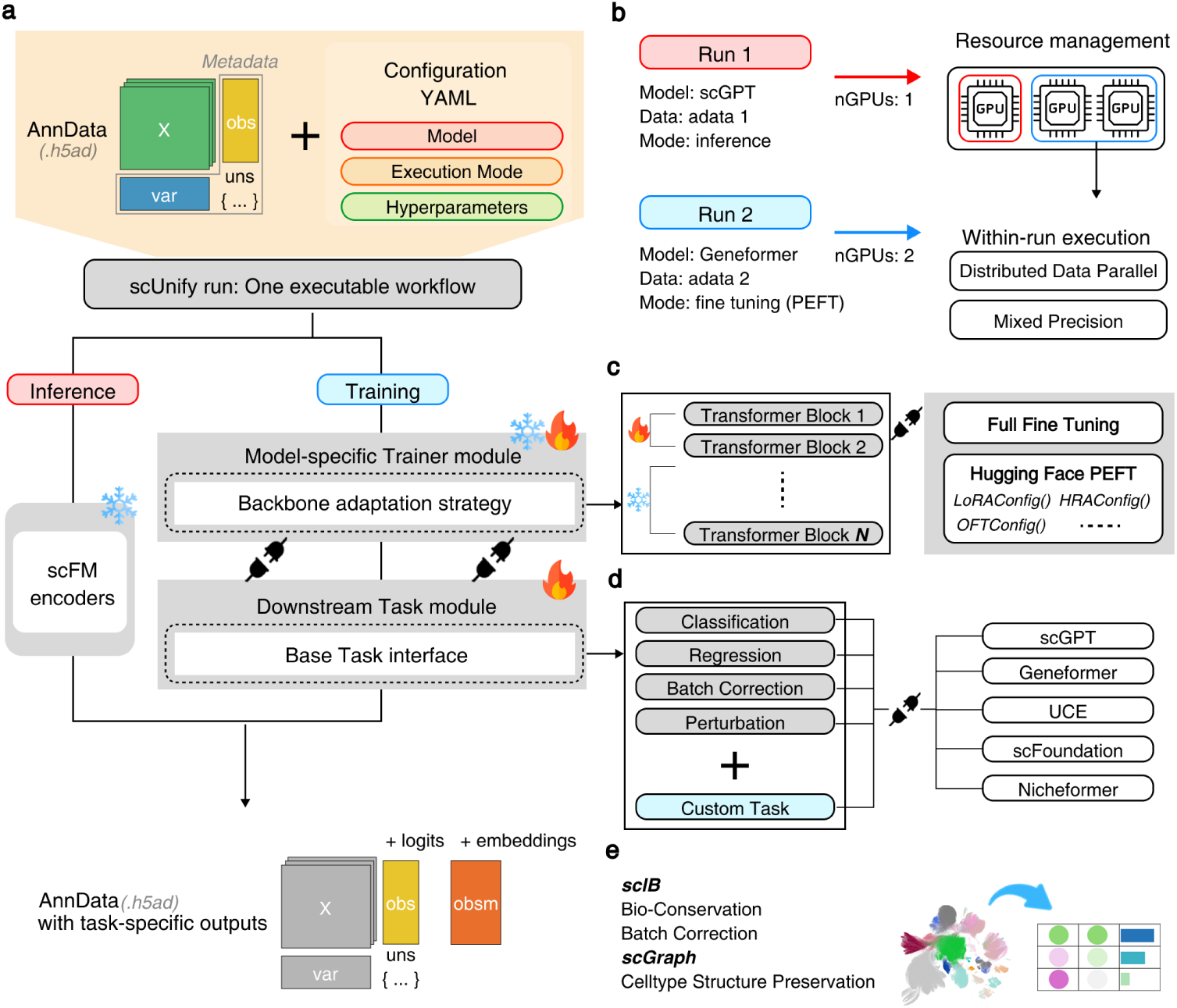
Graphical overview of the scUnify framework. (a) Overall workflow of scUnify. An AnnData object and a configuration file define an scUnify run, which performs inference or training through model-specific processing and returns cell embeddings and task-specific outputs in AnnData. (b) The scUnify core system. Multiple runs are scheduled across available computational resources, with mixed-precision computation and Distributed Data Parallel supported within each run. (c) Adaptation strategies. Backbone adaptation is configured independently of the downstream task module to support Full-FT and multiple Hugging Face PEFT methods. (d) Extension of trainable downstream tasks across scFM backbones. Built-in and custom downstream task modules are implemented through a common base task interface, allowing the same task-learning logic to be connected to and reused across compatible model-specific trainers. (e) Integration of cell-embedding evaluation metrics in scUnify.

Comparing multiple models and execution conditions for the same dataset and analytical objective requires repeated management of individual analyses and computational resources. scUnify defines one analysis execution as an scUnify run comprising an AnnData object and a configuration file, which specifies the model, execution mode, downstream task, adaptation strategy, and hyperparameters. When multiple runs are submitted, the Ray-based core system[20] assigns each run to the appropriate model environment and available CPUs and GPUs, and manages queueing or concurrent execution according to user-specified resource requirements and resource availability (Figure 1b). Within an individual run, Hugging Face Accelerate[21] can apply mixed-precision computation and multi-GPU execution using Distributed Data Parallel (DDP). Researchers can therefore manage runs within the same execution system while reducing the repeated work of separately coordinating environments and resource allocations even when the number of candidate scFMs and conditions increases.

The training mode of scUnify is organized around three independently configurable components: the model-specific trainer, the adaptation strategy, and the downstream task module (Figure 1a, c, d). The model-specific trainer handles model-specific preprocessing, input construction, pretrained-backbone loading, and the training forward pass for the five models currently included in scUnify: scGPT, Geneformer, UCE, scFoundation, and Nicheformer[2–6]. Within each trainer, the transformer backbone is connected to the official Hugging Face PEFT library[22] through model-specific target mappings. It allows researchers to change the backbone training scope through probing, partial fine-tuning, Full-FT, or supported PEFT methods without modifying model specific trainer or the task-learning logic (Figure 1c). Meanwhile, the downstream task module uses a common base task interface(Figure 1d). This interface implements the prediction architecture, objective, evaluation, and output logic required for a trainable task, and can connect that task to an existing model-specific trainer. Based on this structure, scUnify provides classification, regression, batch correction, and perturbation prediction as built-in downstream tasks derived from the trainable workflows reported for the five scFMs (Supplementary Table S2). Researchers can therefore reuse a supported built-in task across compatible backbones or even extend a new trainable task through the same interface, while independently applying and comparing adaptation strategies without reimplementing every model–task–adaptation combination.

scUnify provides a separate inference mode for generating cell representations from a frozen backbone without task-specific training. The model-specific inferencer applies model-specific preprocessing and input construction to the input AnnData and performs a forward pass through the frozen pretrained backbone. The resulting cell embeddings are stored in the appropriate AnnData field. Those embeddings can be evaluated through the public scUnify API, scunify.evaluation.Evaluator (Figure 1e). The Evaluator includes scIB for assessing biological conservation and batch correction[18] and scGraph for assessing celltype structure preservation[19]. The evaluation results and AnnData can then be used in subsequent Scanpy and scverse analyses. Researchers can thus compare cell representations and carry them into downstream analyses without separately converting embedding outputs or reimplementing evaluation procedures for each model.

### 2.2 scUnify reproduces evaluated model outputs and characterizes model-specific computational behavior

To assess the reliability and practical applicability of the common execution workflow, we examined several quantitative index: numerical consistency of cell embeddings with the original implementations, training behavior during continual learning, whether end-to-end inference runtime increased relative to the original workflows under the evaluated conditions, and whether model-specific resource trade-offs could be compared within the same workflow. We also varied numerical precision, batch size, and the number of GPUs allocated to a run and analyzed the outputs and resource use of inference and training. In inference mode, we compared cell embeddings generated by scUnify and the original implementations. In training mode, we performed continual learning using each model’s pretraining objective and tracked the loss trajectories; Geneformer and Nicheformer, whose original repositories provided comparable continual learning workflows, were also compared directly with their original workflows. The mean cell-wise cosine similarity was 0.9926–1.0000 across all model–precision conditions (Figure 2a). Across precisions, scUnify losses generally followed similar decreasing trajectories within each model, and the directly compared training-loss trajectories closely matched those of the original workflows (Figure 2d). However, the FP16 loss for scFoundation became unstable late in training, whereas its BF16 loss decreased stably. UCE also exhibited precision-dependent optimization behavior, with nearly overlapping FP16 and BF16 loss trajectories but a distinct FP32 trajectory. These results indicate that scUnify generated cell embeddings that were numerically consistent with the original implementations across all five models and maintained training fidelity in the directly comparable continual learning workflows, while numerical precision needed to be selected with stability in mind.

**Fig. 2.**
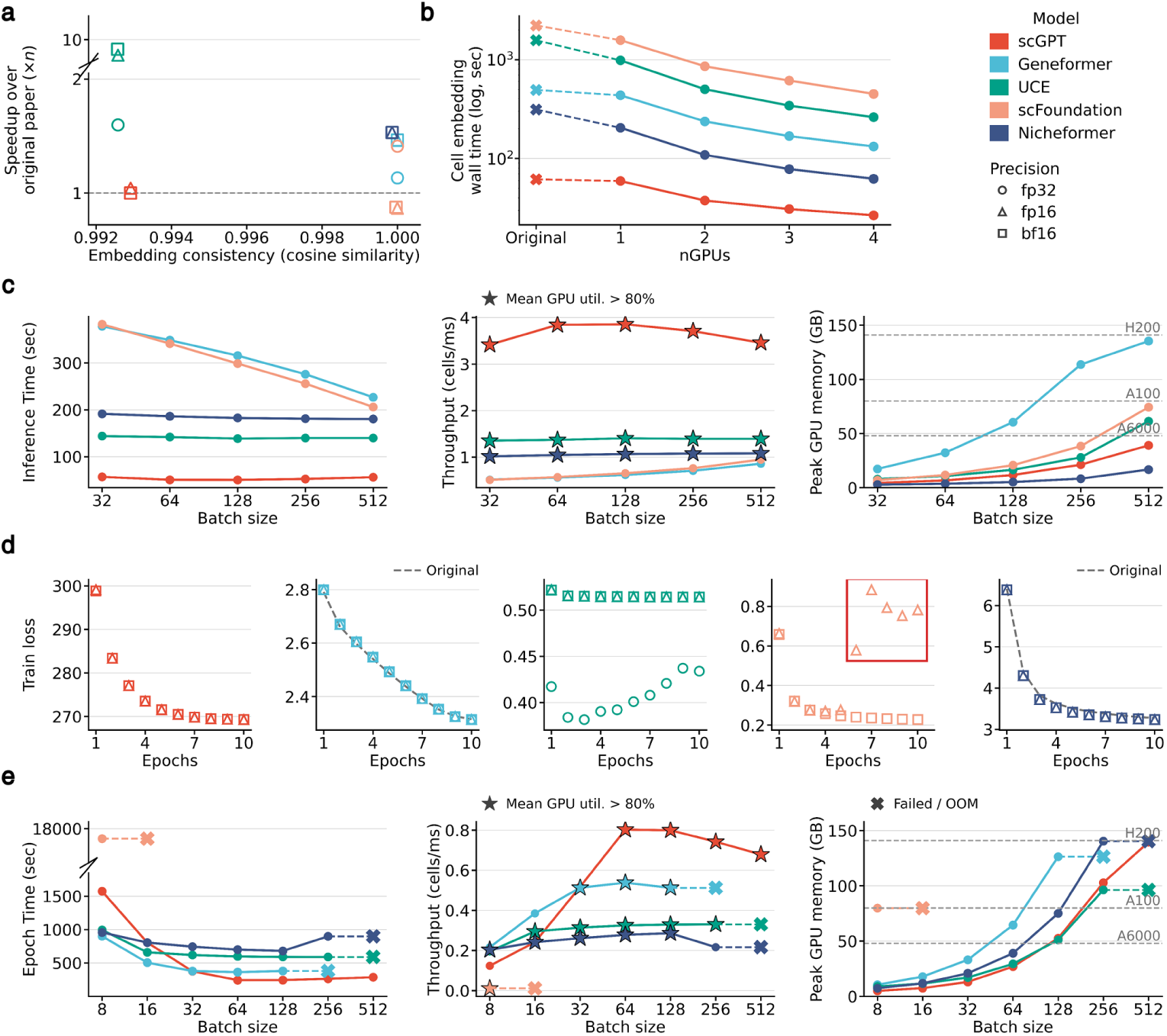
Resource profiling of scUnify inference and training workflows. (a) Precision sweep for cell-embedding inference. Each point shows the mean cosine similarity between scUnify and the original models’ cell embeddings, together with the speedup over the original workflow. Colors indicate models and marker shapes indicate numerical precision. (b) Multi-GPU scaling of cell-embedding inference. Original workflows were run on a single GPU, whereas scUnify was evaluated with one to four GPUs. Wall time includes data loading and model execution. (c) Inference batch size sweep showing inference time, throughput, and peak GPU memory. Stars indicate settings with mean GPU utilization above 80%. (d) Training loss trajectories for the default pretraining-style objective across precision settings. Original-code loss curves are shown as gray dashed lines where available. Precision markers were horizontally offset for visibility. (e) Training batch size sweep showing second-epoch wall time, throughput, and peak GPU memory. X indicate the first failed or out-of-memory batch size.

Next, we compared end-to-end cell-embedding inference runtime between scUnify and the original workflows at the numerical precision used by each original implementation. Under these evaluated conditions, all five models had shorter wall times with scUnify than with their original workflows, with speedups of 1.04–1.60× (Figure 2a). Mixed precision further reduced runtime for some models. Under FP16 and BF16, UCE showed speedups of 9.61–9.76× relative to its original FP32 workflow. scFoundation achieved a 1.41× speedup in the matching FP32 comparison, whereas runtime increased under mixed precision with the same batch size of 1 as the original implementation. With multi-GPU execution using Distributed Data Parallel (DDP), increasing the allocation from 1 GPU to 4 GPUs reduced wall time for every model, with model-specific speedups of 2.22–3.75× (Figure 2b). For subsequent experiments, we jointly considered numerical consistency, training stability, and runtime and used FP16 for scGPT and BF16 for the other four models.

Finally, we fixed the selected numerical precision and compared model-specific resource trade-offs on a single GPU across inference batch sizes of 32–512 and training batch sizes of 8–512. Because inference does not update parameters, its runtime for each model was shorter than the training epoch time, and throughput generally improved as batch size increased (Figure 2c). However, inference throughput across successful conditions varied by model and batch size over a range of 0.51–3.86 cells/ms, and models also differed in where throughput plateaued or began to decline and in how peak GPU memory increased. For training profiling, each backbone was coupled to and trained with the prediction or reconstruction module required by its model-specific pretraining objective. Across successful conditions that included gradient computation and parameter updates, training throughput was 0.011–0.802 cells/ms, and the batch size that maximized throughput and the point of execution failure or out-of-memory differed among models (Figure 2e). For example, scFoundation jointly trained a decoder that reconstructed expression for 19,264 genes and exhibited longer epoch times and higher memory requirements than the other models. scUnify records wall time, throughput, mean GPU utilization, and peak GPU memory consistently for each run, allowing users to compare the computational costs and resource trade-offs associated with different backbones and model-specific learning structures within the same workflow. Together, scUnify maintained numerical consistency with the original implementations across all five models and training fidelity in the directly comparable continual learning workflows, did not increase end-to-end inference runtime relative to the original workflows under the evaluated conditions, and revealed resource trade-offs among scFMs on a common basis.

### 2.3 scUnify enables systematic comparison of scFM backbones and adaptation strategies across downstream tasks

scUnify separates the model-specific trainer, the downstream task module and the adaptation strategy, allowing different combinations of the scFM backbone-downstream task-adaptation strategy to be executed and compared within an integrated workflow. To evaluate whether this separation could extend model-native downstream workflows with additional adaptation strategies without reimplementing task-specific learning logic, we configured trainable workflows reported across the five scFMs as built-in downstream tasks(Supplementary Table S2, Figure 3). Although their prediction targets differed, the original studies of the four scFMs other than UCE each reported a trainable cell-level classification workflow. We retained their model-native workflows(architectures and objectives) and extended celltype classification to UCE, allowing all five models to be evaluated in a common downstream context. Additionally, we included niche-density regression for Nicheformer, batch correction and perturbation prediction for scGPT, and perturbation prediction for scFoundation. For each downstream task, the model-specific task architecture was kept fixed while adaptation strategies were independently applied to the corresponding backbone.

**Fig. 3.**
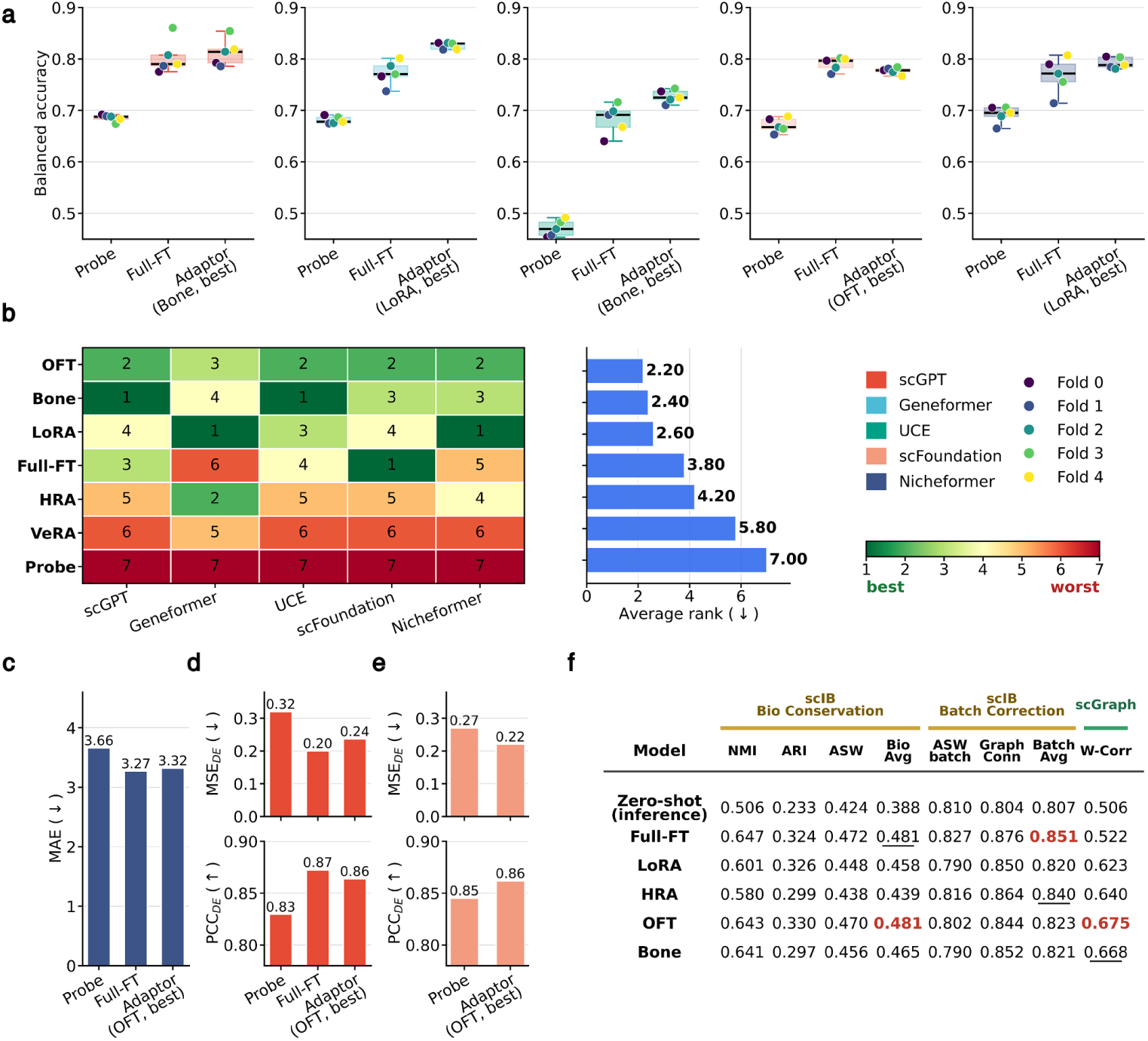
Performance of trainable downstream tasks across scFM backbones. (a) Balanced accuracy of celltype classification task with model-native workflow. (b) Adaptation strategies ranked within each backbone (left) and by average rank across backbones (right). (c) MAE of Nicheformer regression task. (d-e) Perturbation tasks for scGPT and scFoundation. (f) scIB bio-conservation, scIB batch correction, and scGraph Weighted Correlation of scGPT Batch correction task. The best value is red and the second is underlined.

We first evaluated this design by applying all five scFMS to NSCLC(non-small cell lung cancer) celltype classification (Figure 3a-b). With each model-specific classification architecture held fixed, we compared probing and full fine-tuning (Full-FT) with five parameter-efficient fine-tuning (PEFT) methods: low-rank adaptation (LoRA), Householder reflection adaptation (HRA), orthogonal fine-tuning (OFT), vector-based random matrix adaptation (VeRA), and block affine adaptation (Bone). Full-FT achieved higher balanced accuracy than probing for every backbone, indicating that training the backbone was generally more effective than using a frozen pretrained representation for this classification task. For four backbones other than scFoundation, at least one PEFT method outperformed Full-FT, showing that adapters independently connected to the backbone while retaining the model-native classification architecture could also improve downstream performance. When the seven adaptation strategies were ranked within each backbone, OFT and Bone achieved the lowest average ranks of 2.20 and 2.40, respectively, whereas probing ranked last for every backbone (Figure 3b). However, OFT, which achieved the best average rank, was not the top-performing condition for any individual backbone. Instead, the top-performing condition was Bone for scGPT and UCE, LoRA for Geneformer and Nicheformer, and Full-FT for scFoundation. Thus, although Full-FT and PEFT were generally more effective than probing, an adaptation strategy with a strong average rank was not necessarily optimal for an individual backbone.

When the evaluation was extended beyond classification to other built-in downstream tasks, no single adaptation strategy performed consistently best. Full-FT achieved the lowest MAE in Nicheformer niche-density regression (Figure 3c) and both the lowest *MSE_DE_* and the highest *PCC_DE_* in scGPT perturbation prediction (Figure 3d), whereas OFT performed best among the evaluated conditions for scFoundation perturbation prediction (Figure 3e). For scGPT batch correction, we used scIB and scGraph integrated into scUnify to evaluate the zero-shot cell representation alongside Full-FT and PEFT conditions. Although the trained conditions generally improved upon zero-shot inference, the strongest adaptation strategy differed across evaluation aspects. OFT and Full-FT produced similar average biological-conservation scores of 0.481; OFT achieved the highest scGraph weighted correlation of 0.675, whereas Full-FT achieved the highest average batch-correction score of 0.851 (Figure 3f). Taken together, adapters independently applied to each model-native downstream architecture were competitive with or outperformed Full-FT under several conditions, but their relative benefits and the top-performing adaptation strategy varied across backbones and downstream tasks. Determining a suitable condition for a target dataset and downstream task therefore requires direct execution and comparison of Full-FT and PEFT conditions across candidate backbones, which scUnify enables within a consistent training and evaluation workflow.

### 2.4 scUnify integrates a custom task with multiple scFM backbones

To evaluate whether scUnify could implement a new trainable task once and extend it across multiple scFM backbones and adaptation strategies, we implemented scMILD-based multiple instance learning (MIL)[23] for sample-level COVID-19 infection prediction as a custom downstream task module. This task required a MIL-specific Dataset and DataLoader that organized cells into sample-level bags, an alternating procedure for training the sample and cell branches, and separate optimization of the backbone and the two branches (Figure 4a, b). To accommodate the GPU memory requirements of scFM backbones, we introduced a cell-sampling step into the scMILD workflow that limited the number of cells processed per sample while approximately preserving its celltype composition. The same sampling procedure was applied to the autoencoder-based scMILD baseline and the scUnify variants, in which the original MLP encoder was replaced with a Geneformer, scGPT, scFoundation, UCE, or Nicheformer backbone. The resulting MIL workflow was trained and compared across multiple combinations of scFM backbones and adaptation strategies using the same task architecture and common training settings.

**Fig. 4.**
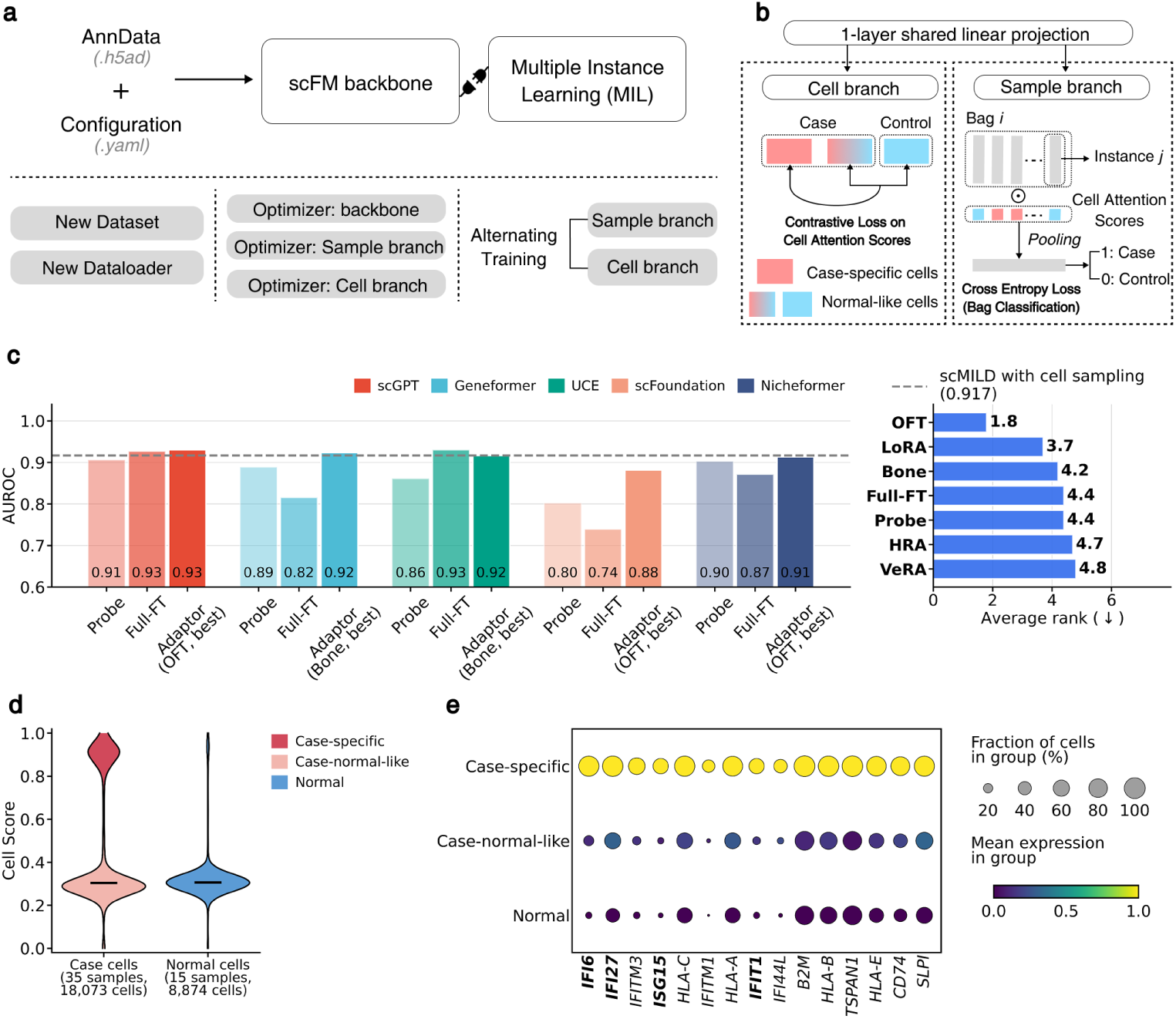
scUnify custom MIL task. (a) scUnify custom downstream task, and its custom modules and training strategies. (b) scMILD architecture overview. (c) AUROC of the MIL task (left) and average rank of adaptation strategies (right). (d) Cell-score distribution in overall samples and colors indicate cell groups. (e) Differentially expressed genes between case-specific cells and case-normal-like/normal cells. Dot size indicates the fraction of cells expressing each gene, and color indicates the gene-wise min–max-scaled mean expression.

We compared sample-level AUROC across the five backbones under probing, full fine-tuning (Full-FT), and five PEFT methods: LoRA, HRA, OFT, VeRA, and Bone (Figure 4c). scGPT and Nicheformer performed strongly under multiple adaptation conditions, whereas Geneformer improved substantially with Bone compared with Full-FT. When the ranks of the seven adaptation strategies were averaged across backbones, OFT achieved the best average rank, although the top-performing PEFT method for individual backbones differed between OFT and Bone. The autoencoder-based scMILD baseline achieved a mean AUROC of 0.9167 when evaluated using the same cell-sampling procedure as the scUnify experiments; this baseline was exceeded by scGPT with Full-FT or OFT, Geneformer with Bone, and UCE with Full-FT, showing that different backbones could reach performance above the baseline through either Full-FT or PEFT. To compare the relative performance of the backbones observed in MIL with that in another downstream context, we also repeated NSCLC celltype classification using the one-layer classifier and a standardized training setting across all five backbones instead of their model-native classification architectures (Supplementary Fig. S1). Under this setting, the absolute performance and rankings of some adaptation strategies changed relatively to the model native workflows in Figure 3a. Nevertheless, when the best-performing condition for each backbone was compared, UCE remained below the other four models, consistent with its relative position in the model-native results. In the MIL task, by contrast, UCE ranked among the stronger backbones whereas scFoundation shifted to a lower relative position, showing that their relative performance differed across downstream contexts.

Because scMILD produces attention-derived cell scores alongside sample-level predictions, the cell populations contributing to predictions from a selected model can be analyzed further. We used cell scores from a representative high-performing scGPT–OFT condition to divide cells into case-specific, case-normal-like, and normal groups (Figure 4d). Comparing the case-specific group with the other two groups in Ciliated cells showed higher mean expression and fractions of expressing cells for interferon-response genes, including *IFI6*, *IFI27*, *ISG15*, and *IFIT1* (Figure 4e). This observation indicates that, in the representative scGPT–OFT condition, the cell-level expression pattern was consistent with the interferon-response pattern reported in the original scMILD study[23]. These results show that changing the scFM backbone within the same custom downstream task structure can improve sample-level predictive performance while retaining the task’s cell-level biological interpretability. More broadly, in an scFM ecosystem where relative model performance varies across downstream contexts, these findings support the role of scUnify as a framework for implementing a new trainable task once, reusing it across multiple scFM backbones with independently application, and comparing adaptation strategies within a common workflow.

### 2.5 scUnify provides supporting utilities for researchers

To support configuration preparation, scUnify provides assistant skills that enable external coding assistants such as Claude Code and Codex to consult its public documentation (Figure 5a). Claude Code uses .claude/skills/scunify/SKILL.md, whereas Codex uses .agents/skills/scunify/SKILL.md; both skills direct the assistants to the public documentation under docs/, which describes installation, configuration, supported models, execution modes, and downstream tasks. Based on these resources and a user’s request, a coding assistant can help prepare a configuration file, execution script, or notebook and answer questions about using scUnify. The prompt in Figure 5a is an illustrative example of this workflow.

**Fig. 5.**
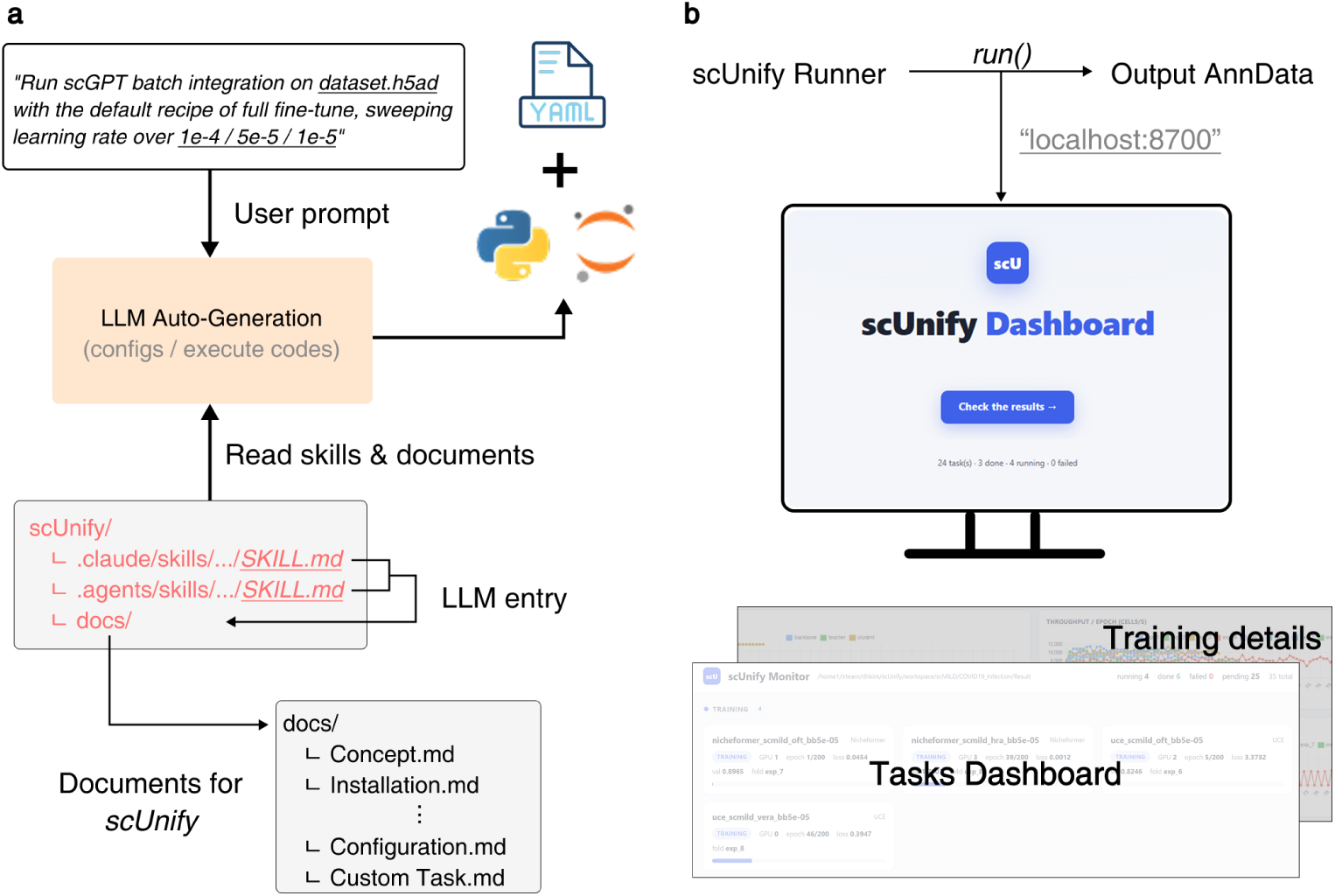
Supporting utilities of scUnify. (a) Schematic of usage with an LLM assistant, in which external coding assistants read scUnify’s documentation through their entry points (Claude Code via.claude/skills/, Codex via .agents/skills/). (b) The scUnify dashboard, showing submitted task status and per task training progress during execution.

scUnify also provides a web dashboard that can be used with ScUnifyRunner to inspect run status and training progress during long-running executions that include multiple conditions (Figure 5b). The dashboard summarizes the pending, training, done, and failed states of multiple runs in a single view and displays records such as epoch, loss, validation results, and throughput for runs in progress. This view allows users to inspect the status of multiple runs and the training progress of each run during execution.

## 3 Discussion

In this study, we developed scUnify to connect scFM execution, downstream task learning, and adaptation across distinct software environments and implementation procedures within a common workflow. Whereas existing scFM integration frameworks have primarily focused on providing common execution and benchmarking infrastructure or extending trainable downstream task interfaces[13–16], scUnify differs by separating the model-specific trainer, downstream task module, and adaptation strategy into independent components and supporting their combinations within a unified execution system. In this design, the model-specific trainer preserves the preprocessing, tokenization, and forward procedure of each scFM, the downstream task module defines the task-specific architecture and objective function, and the adaptation strategy independently controls the training scope of the backbone without changing the task-learning logic. Existing downstream tasks can therefore be executed across multiple model and adaptation conditions, while a new trainable task can be implemented once and reused across supported backbones and adaptation strategies. Based on this structure, we implemented five built-in downstream workflows spanning classification, regression, batch correction, and model-specific perturbation prediction, following the workflows reported in the original scFM studies. We further implemented the complex MIL-specific data preparation and dual-branch learning procedure as a single custom downstream task module and demonstrated its extension across five scFM backbones and multiple adaptation strategies.

Our results illustrate the practical need for this combinatorial design, as the relative performance of both backbones and adaptation strategies varied across downstream contexts. In celltype classification using the same one-layer classifier and standardized training setting, UCE performed below the other backbones, whereas scFoundation ranked relatively high. In the custom MIL task, however, UCE showed relatively strong performance, while scFoundation moved to a lower position. These contrasting results indicate that the relative performance of a backbone observed for one downstream task and dataset cannot be directly generalized to another analytical context. Broad tendencies within individual models also did not always identify the best adaptation strategy for a specific condition. Although Full-FT and PEFT generally outperformed probing, and OFT and Bone achieved comparatively strong average ranks across multiple comparisons, Full-FT remained the best strategy for some tasks, while the top-performing PEFT method varied among LoRA, OFT, and Bone depending on the backbone and downstream task. These observations are consistent with previous benchmarking studies showing that the relative performance of scFMs and adaptation strategies can vary with the dataset and downstream context[7, 8, 12], further indicating that the superiority of a particular model or adaptation strategy cannot be generalized from a single benchmark. Researchers therefore need an integrated framework that allows candidate model–task–adaptation combinations to be examined directly through a consistent execution and evaluation process for their target dataset and downstream task, and scUnify enables such comparisons within a common workflow.

Building on this implementation design, scUnify also provides usability features that support researchers throughout practical analysis workflows. First, its core system schedules multiple runs across available CPUs and GPUs and supports mixed-precision computation and Distributed Data Parallel within each run. Second, cell representations generated through training or inference can be evaluated within the same workflow using the scUnify Evaluator, which integrates scIB and scGraph. Third, assistant skills guide external LLM-based coding assistants to public documentation during configuration preparation, while the runtime dashboard allows researchers to monitor the status and training progress of multiple runs during execution. These usability features were integrated without compromising fidelity to the original workflows in inference and training, and scUnify achieved faster execution under some evaluated conditions.

Meanwhile, adding a new scFM to scUnify still requires implementing a model-specific trainer that reflects its preprocessing and input construction, backbone loading, forward procedure, and software dependencies, followed by validation against the original implementation. Once added, however, the model-specific trainer can be reused with existing downstream task modules and adaptation strategies, reducing the repetitive model-specific implementation required for subsequent task and adaptation experiments. Future work will expand the supported backbones and downstream tasks, consolidate recurring trainer elements into common functionality, and systematize validation procedures for new model-specific implementations. Although coding assistants can use the assistant skills to consult the tutorial documentation when preparing individual configuration files, each scUnify run is currently defined by a separate configuration file. Consequently, as the number of model–task–adaptation and hyperparameter conditions grows, so does the number of configuration files that must be managed separately. Because the scUnify core system is built on Ray, future work will integrate Ray Tune so that multiple experimental conditions can be defined and explored from a single configuration.

## 4 Conclusions

We developed scUnify, a framework that connects scFMs separated by different implementations and execution environments so that they can be executed, compared, and extended within a common analytical workflow. Our evaluation showed that scUnify retained the reliability of integrated execution while enabling direct comparison of model–adaptation combinations in the target context and extension of a complex downstream task, once implemented, across multiple supported scFMs. scUnify connects the execution of heterogeneous scFMs and provides a common basis for researchers to compare models and adaptation strategies on their own datasets and for their analytical objectives, and to extend new trainable downstream tasks across multiple backbones.

## 5 Methods

### 5.1 Implementation details

scUnify establishes a common workflow spanning data input, model execution, downstream task training, and output storage for five single-cell RNA-sequencing foundation models (scFMs): Geneformer[3], scGPT[2], scFoundation[4], Nicheformer[6], and UCE[5]. The implementation comprises task definition and configuration, a Ray-based core system, model-specific trainers, downstream task modules, and adaptation strategies. The relationships among these components and classes are shown in Supplementary Fig. S2, and the runtime organization of the scUnify core system is shown in Supplementary Fig. S3.

**Task definition and configuration.** scUnify defines an analysis execution as an scUnify run comprising an AnnData file (.h5ad) and a YAML configuration file (.yaml). AnnData contains the expression matrix and metadata, whereas the configuration file specifies the model, execution mode, downstream task, adaptation strategy, and execution conditions. ScUnifyConfig interprets these inputs as a run configuration, and ScUnifyRunner executes one or more runs according to their configurations. The resulting cell embeddings and task-specific outputs are stored in the obs, obsm, or uns fields of AnnData, allowing inputs, predictions, and embeddings to be passed within the same data object to subsequent Scanpy[24] and scverse[17] analyses.

**Inference and training modes.** Inference and training use separate execution paths. In inference mode, a model-specific inferencer applies model-specific preprocessing and input construction and generates cell embeddings through forward passes of a frozen pretrained encoder. In training mode, a model-specific trainer is combined with a downstream task module, trainable parameters are selected according to the chosen adaptation strategy, and the model is trained using the task objective. After training, predictions and cell embeddings are extracted from the selected checkpoint and stored together with fold-specific AnnData objects, training histories, and summary artifacts.

**scUnify core system.** When multiple runs are submitted to ScUnifyRunner, the Ray-based core system[20] selects the model-specific Conda environment specified in each configuration through runtime env and manages resource allocation, queueing, and concurrent execution according to the CPU and GPU requests of each run and the available resources. Runs using the same model environment and dataset share a Ray Object Store reference to the AnnData loaded by DataLoaderActor. Each run is executed using Ray TorchTrainer, within which Hugging Face Accelerate[21] handles device placement, mixed-precision computation, and multi-GPU execution using Distributed Data Parallel (DDP). ProgressActor records inference-batch progress and the fold, epoch, loss, validation result, and throughput of training, and a web dashboard implemented with FastAPI [25] displays run states and training progress.

**Separation of model-specific trainers and downstream task modules.** In training mode, model-specific trainers and downstream task modules are implemented as independent components. A model-specific trainer constructs backbone inputs from AnnData, loads pretrained weights, and handles the training forward procedure and extraction of cell representations. Built-in and user-defined custom downstream tasks are implemented as task interfaces that subclass TaskMixin and follow a common interface for the prediction architecture receiving backbone outputs, objective, evaluation procedure, and AnnData output logic. A custom task subclass defines a task-specific prediction module, loss and output logic. At runtime, scUnify dynamically composes the task interface and model-specific trainer selected in the configuration into a single composed trainer class through multiple inheritance, allowing the same task to be attached to supported scFM backbones.

**Backbone adaptation strategies.** The adaptation configuration is separated from downstream task learning and specifies the trainable backbone layers and, when PEFT is used, the adapter configuration. Probing freezes the backbone and trains only downstream task parameters; partial fine-tuning trains only selected Transformer layers; and full fine-tuning (Full-FT) trains the entire backbone and task-specific parameters. PEFT uses the official Hugging Face PEFT integration[22], with compatible Transformer encoder modules in each scFM mapped to Hugging Face target modules. Users can select PEFT configurations that are available in the installed Hugging Face PEFT package and compatible with the chosen backbone. Model-specific attention transformations and target-module mappings are provided in Model-specific implementation details in the Supplementary Methods S1.

**Assistant skills and public documentation.** The scUnify repository provides assistant skills that direct external coding assistants to the user documentation and tutorials in the public docs/ directory (Fig. 5a). The entry point for Claude Code is .claude/skills/scunify/SKILL.md, and that for Codex is.agents/skills/scunify/SKILL.md; both files route requests concerning installation, configuration, inference and training, built-in and custom downstream tasks, adaptation, evaluation, outputs, and monitoring to the corresponding docs/ pages. Based on these materials and a user’s request, a coding assistant can support the preparation of configuration files, execution scripts, or notebooks.

### 5.2 Dataset and preprocessing

Six single-cell datasets were used for resource profiling and downstream task evaluation. The CellTypist Kidney reference was used for inference and training resource profiling of the five scFMs[26]. A NSCLC(non-small-cell lung cancer) tumor-infiltrating T-cell dataset[27] was used for celltype classification with the five backbones, the COVID-batch subset[2] for scGPT batch correction, and the Norman CRISPRa Perturb-seq dataset[28] for perturbation prediction with scGPT and scFoundation; the preprocessed datasets provided by scPEFT[12] were used for these three analyses. A 10x Xenium human lung dataset was used for Nicheformer neighborhood-density regression[6], and a COVID-19 nasal-swab dataset[29] was used for the custom multiple-instance learning (MIL) task. Each dataset was provided in AnnData format with an expression matrix, task labels, conditions, sample identifiers, and split assignments. Dataset sources, sizes, preprocessing, and splits are provided in Supplementary Table S3.

### 5.3 Experimental design

The evaluation comprised resource profiling of inference and training using model-specific pretraining objectives across the five scFMs, built-in downstream task experiments, and a custom MIL experiment. The built-in experiments included celltype classification across all five backbones, Nicheformer regression, scGPT batch correction and perturbation prediction, and scFoundation perturbation prediction (Fig. 3). For each model–task pair, the downstream architecture, objective, and task-specific training settings followed the corresponding original workflow where available[2–6]. The custom MIL experiment applied the same scMILD-based downstream task module[23] to all five backbones (Fig. 4). To minimize differences arising from downstream task design, the task architecture and training parameters were held constant across backbones, while the learning rate differed only between adaptation conditions. For the same reason, we additionally performed standardized celltype classification by connecting all five backbones to the same one-layer classifier and using common training settings other than learning rate (Supplementary Fig. S1). Across the downstream experiments, we compared probing, full fine tuning (Full-FT), and five PEFT methods: low-rank adaptation (LoRA)[30], Householder reflection adaptation (HRA)[31], orthogonal fine-tuning (OFT)[32], vector-based random matrix adaptation (VeRA)[33], and block affine adaptation (Bone)[34]. Detailed model- and task-specific hyperparameters are provided in Supplementary Table S10.

### 5.4 Resource profiling

Inference and training resource profiling used the CellTypist Kidney dataset as a common reference and measured wall time, throughput, mean GPU utilization, and peak GPU memory. Inference profiling evaluated numerical consistency, end-to-end wall time, batch size response, and multi-GPU scaling (Fig. 2). Numerical consistency was calculated as the mean cell-wise cosine similarity between cell embeddings generated for the same cells by scUnify and the original implementation, 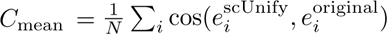, and was compared across supported numerical-precision conditions, including the original condition. End-to-end wall time was defined as the interval from data loading through cell-embedding generation, and speedup over the original implementation was calculated as *S*_inference_ = *T*_original_*/T*_scUnify_. Throughput in the batch size sweep was calculated from model-forward inference time and reported in cells/ms. Multi-GPU scaling was evaluated by comparing DDP inference wall time while varying the number of GPUs assigned to one run. Meanwhile, training resource profiling used the model-specific pretraining objective of each backbone for continual learning on the Kidney dataset. Direct comparisons of training-loss trajectories with the original workflows were performed for Geneformer and Nicheformer because their original repositories provided comparable continual learning workflows[3, 6]. Loss trajectories were compared across numerical precisions, and per-epoch wall time, training throughput (cells/ms), mean GPU utilization, peak GPU memory, and execution failure or out-of-memory status were recorded while varying batch size. The architecture and loss for each objective are described in the Supplementary Methods S2.

### 5.5 Downstream Tasks

The built-in downstream tasks in scUnify comprised celltype classification applied to all five scFM backbones and the tasks used for individual models in their original studies: Nicheformer regression, scGPT batch correction, and perturbation prediction with scGPT and scFoundation (Fig. 1d). For all tasks, the input expression matrix in AnnData was denoted by *X* = (*x_ij_*) ∈ R*^N^*^×^*^G^*, where *N* and *G* denote the numbers of cells and genes, respectively. Each backbone applied model-specific preprocessing and input construction to generate hidden representations and cell representations, which the downstream task module passed to a task-specific prediction architecture and objective. Graphical overviews of the downstream architectures and notation are shown in Supplementary Figs. S4-S6.

**Evaluation metrics.** celltype classification was evaluated using balanced accuracy, Nicheformer regression using mean absolute error (MAE), perturbation prediction using the mean squared error and Pearson correlation coefficient for the top differentially expressed genes between perturbed and control cells (MSE_DE_ and PCC_DE_)[2, 4, 35], and the custom MIL task using sample-level Area Under the Receiver Operating Characteristic curve (AUROC). Batch-corrected cell representations were evaluated using the biological-conservation and batch-correction scores from scIB[18] and the celltype structure-preservation metrics from scGraph[19]. Balanced accuracy, MAE, MSE, and AUROC were calculated using scikit-learn [36], and Pearson correlation coefficients were calculated using SciPy [37]. Detailed metric definitions are provided in Supplementary Table S4.

**Celltype classification and Nicheformer regression.** T cells in the NSCLC dataset were classified into *K* = 14 subtypes. Geneformer, scGPT, scFoundation, and Nicheformer each used the downstream classification architecture from the corresponding original implementation. Because the original UCE study focused on zero-shot evaluation using frozen representations and did not provide a trainable classification head, a one-layer MLP classifier was used for UCE. The observed label of cell *i* was denoted by *y_i_* ∈ {1*, …, K*}, and the probability assigned by the classifier to class *c* by *p_ic_*; multiclass cross-entropy was minimized.

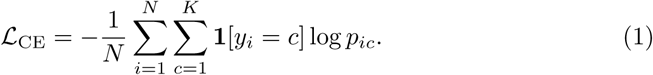

Nicheformer regression predicted the number of neighboring cells located within 25 *µ*m of each cell in the Xenium Lung dataset. The regression head used the same architecture as the Nicheformer classification head and produced a scalar output with *K* = 1. The target and prediction for cell *i* were denoted by *y_i_* and *y*^*_i_*, respectively, and mean squared error was used as the training objective.

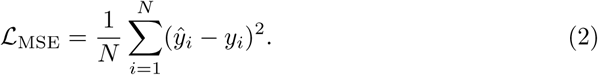

**Batch correction.** For scGPT batch correction, domain-specific batch normalization (DSBN) conditioned on the batch label in the COVID-batch dataset was applied to input representations, and encoder outputs and batch embeddings were passed to gene-expression prediction (GEP), masked value prediction from the cell representation (MVC), domain-adversarial batch classification (DAB), and elastic cell similarity (ECS) objectives (Supplementary Fig. S5a)[2]. The overall objective was

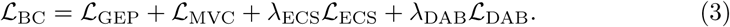

GEP used the hidden representations from the final encoder layer and a batch embedding to predict expression and the probability of zero expression for masked gene tokens. Denoting the GEP expression prediction and zero-expression probability at masked positions by 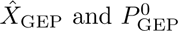, respectively, its objective was

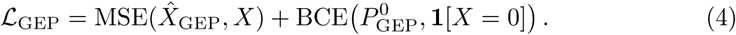

MVC combined the cell representation and batch embedding with the gene-token embeddings formed before input to predict expression and the probability of zero expression for masked gene tokens. Denoting the MVC outputs by 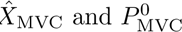, its objective was

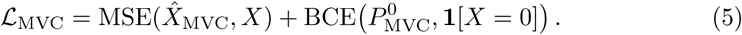

DAB passed the cell representation through a gradient-reversal layer to an MLP batch discriminator that predicted batch probabilities Π^batch^. For the batch-label vector **b**, the DAB objective was

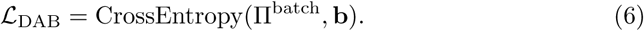

ECS used L2-normalized cell representations *E* to regularize the pairwise cell similarity *S* = *EE*^⊤^ relative to a threshold *τ*.

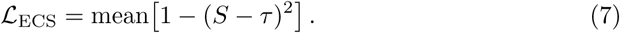

After training, the L2-normalized cell representation was used as the batch-corrected embedding.

**Perturbation prediction.** For perturbation prediction on the Norman CRISPRa Perturb-seq dataset, control expression *X*^ctrl^ and a perturbation condition were used to predict perturbed expression *X*^pert^. For scGPT, gene, expression-value, and perturbation-condition embeddings were summed and passed to the scGPT backbone, and the output hidden representations were passed to coefficient and bias decoders to predict gene-wise *A* and *β*, respectively (Supplementary Fig. S5b). Predicted perturbed expression was calculated as

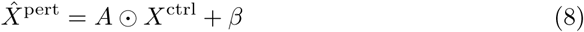

and mean squared error with respect to the observed perturbed expression was minimized.

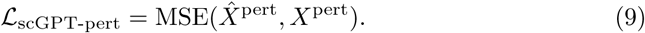

For scFoundation, control expression was passed through the encoder and decoder to generate gene-level representations, which were adjusted using a gene co-expression graph. Learnable embeddings of perturbed genes were adjusted using a gene-ontology graph and summed within each condition to construct a cell-specific perturbation representation. A GEARS[35] decoder combining the two representations predicted gene-wise expression changes Δ*X* (Supplementary Fig. S6).

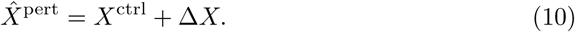

The GEARS objective combined a fourth-power reconstruction error between observed and predicted expression with a penalty for disagreement in perturbation direction relative to the control mean.

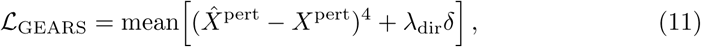

where, denoting mean control expression for each gene by *µ*_ctrl_, *δ* was defined as

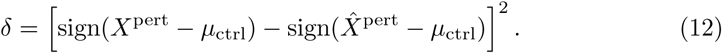

### 5.6 Custom downstream task experiment

To predict sample-level infection status in the COVID-19 infection dataset, scMILD-based dual-branch MIL was implemented as a custom downstream task module[23]. Each donor sample was defined as a bag *s*, each cell *i* within the bag as an instance, and the sample-level label as case or normal. Training, validation, and test assignments were defined at the sample level so that cells from the same sample did not occur in different splits. The autoencoder-based encoder in the original scMILD was replaced with one of the five scFM backbones, and the same downstream task module containing a shared projection, sample branch, and cell branch was applied across backbones and adaptation strategies (Fig. 4a, b).

**Sample branch.** The sample branch transformed the backbone generated cell representation *h_si_* through a shared projection *ϕ* and normalized the score from an attention module *g* within each bag to construct a sample representation *u_s_*. A sample classifier *f* used *u_s_* to predict the case probability *y_s_*.

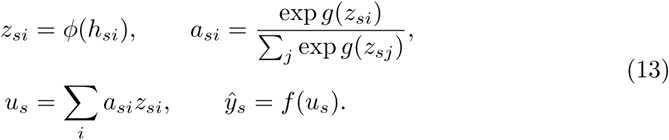

The number of training samples was denoted by *M* and the observed sample label by *y_s_* ∈ {0, 1}, and binary cross-entropy was used as the sample-branch objective.

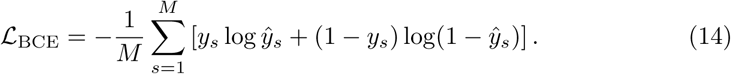

**Cell branch.** The cell branch detached raw attention scores from the sample branch from the gradient, normalized them, and used them as soft pseudo-labels for cells from case samples, whereas cells from normal samples were assigned negative labels. A cell classifier predicted these pseudo-labels using weighted cross-entropy. In addition, a two-component Gaussian mixture model was fitted to cell scores from case samples to distinguish case-specific and case-normal-like groups, and normal cells were included in the negative group together with the case-normal-like group. Orthogonal projection loss (OPL)[38] pulled projected cell representations from the same group together and separated those from different groups. The cell-branch objective was

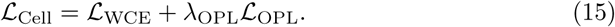

**Training strategy.** The sample and cell branches were trained in alternating epochs at a 1:1 ratio. During a sample-branch epoch, L_BCE_ was used to update the trainable parameters of the shared backbone and projection, whereas L_Cell_ was used during the subsequent cell-branch epoch. To accommodate the GPU memory requirements of scFM backbones, we introduced a cell-sampling procedure for both branches and sampled a fixed number of cells from each sample. At least one cell was randomly selected from each represented celltype, and the remaining cells were allocated according to the within-sample celltype proportions using a multinomial distribution and sampled without replacement.

**Evaluation and Cell score analysis.** Sample-level performance was evaluated using 8-fold cross-validation. AUROC was calculated from the predicted probabilities for the test samples in each fold and reported as the mean across all eight folds. For subsequent cell-level analysis, we selected a median performing fold based on test AUROC among the 8 scGPT with OFT folds. The model obtained from this fold was then applied to all samples in the complete dataset, and the resulting raw cell scores were used for cell-level analysis. Raw cell scores were winsorized at the 1st and 99th percentiles and then min-max normalized. A two-component Gaussian mixture model was fitted to normalized scores from case-sample cells; the higher-scoring component was defined as case-specific and the lower-scoring component as case-normal-like, whereas cells from normal samples were retained as the normal group. Within Ciliated cells, the case-specific group was compared with a reference combining the case-normal-like and normal groups. After total-count normalization to 10,000 counts per cell followed by log1p transformation, the Wilcoxon test implemented in Scanpy sc.tl.rank genes groups was performed[24]. Upregulated genes with adjusted *P <* 0.01 and log fold change *>* 0.25 were ranked by the Wilcoxon score, and the top 15 were displayed in Fig. 4e. Dot size denotes the fraction of cells expressing a gene in each group, and color denotes the group mean expression min–max scaled for each gene across the three groups.

### 5.7 Computational environment and reproducibility

The model-specific Conda environment for each scFM was constructed from the dependencies of its official repository, while Python 3.11.15, Ray 2.54.0, Hugging Face Accelerate 1.13.0, and Hugging Face PEFT 0.14.0 were used across all environments for the common scUnify runtime. Experiments were performed on a server running Ubuntu 22.04.5 LTS and equipped with two Intel Xeon Platinum 8462Y+ processors (64 cores and 128 threads in total), approximately 2.0 TiB of system memory, and four NVIDIA H200 GPUs with approximately 141 GB of memory each. NVIDIA driver version 580.126.16 provided CUDA 13.0 compatibility. Repository revisions, pretrained checkpoints, and model-specific integration details are described in Supplementary Method S1.

OpenAI GPT-5.6 was used to assist with manuscript preparation and English-language editing, while Anthropic Claude Code with Opus 4.8 was used as a software development assistant. All AI-assisted outputs were reviewed and revised by the authors, who take full responsibility for the final software and manuscript.

## Declarations

### Ethics approval and consent to participate

This study involved the secondary analysis of publicly available, de-identified datasets and no direct recruitment of human participants. Ethics approval and informed consent were obtained in the original studies, as reported by the respective authors. No additional ethics approval was required for the present secondary analysis.

### Consent for publication

Not applicable.

### Availability of data and materials

All datasets analyzed in this study were obtained from publicly available sources. The Kidney dataset used for resource profiling was obtained from the CellTypist human kidney reference. The preprocessed NSCLC celltype classification, COVID-19 batch-correction, and Norman perturbation datasets were obtained from the scPEFT Figshare archive (https://doi.org/10.6084/m9.figshare.30763886). The corresponding raw NSCLC and Norman datasets are available from the Gene Expression Omnibus under accession numbers GSE179994 and GSE133344, respectively, whereas the COVID-19 composite comprising 18 batches was originally distributed through the scGPT repository (https://github.com/bowang-lab/scgpt). The Xenium Lung dataset was obtained from SpatialCorpus-110M, and the COVID-19 infection dataset used for the MIL experiment is available from the Broad Institute Single Cell Portal under accession SCP1289. Dataset dimensions, preprocessing procedures, and split definitions are summarized in Supplementary Table S3. The scUnify source code is available at https://github.com/DHKim327/scUnify, with code for reproducing the analyses provided in the Reproduction directory. Documentation for the installation and use of scUnify is available at https://scunify.readthedocs.io/. Code developed as part of scUnify is released under the MIT License, while incorporated third-party model components retain their respective upstream licenses as documented in the repository.

### Authors’ contributions

D.K. conceived and developed the scUnify framework, performed the experiments, and wrote the manuscript. A.H. helped write and edit the English manuscript. K.J. guided the overall direction of the study and edited the manuscript. K.K. supervised the study. All authors reviewed and approved the final manuscript.

## Acknowledgements

Not applicable.

