## Supplementary Information for "scUnify: a unified framework for training and inference across multiple single-cell foundation models"

### Supplementary Methods

#### Method S1. Model-specific implementation details

The five scFMs were implemented from fixed revisions of their official repositories and publicly released pretrained checkpoints. Each model-specific trainer preserves the input construction and forward procedure required by its corresponding model while exposing its cell representation and trainable components through the common scUnify workflow. When a complete continual-learning procedure was unavailable, we implemented the pretraining objective described in the original study and released model configuration. Repository revisions, pretrained checkpoints, and model-specific integration details are provided below.

**scGPT.** The scGPT implementation used revision `cebd6fae` of the official repository at `https://github.com/bowang-lab/scGPT` and its whole-human pretrained checkpoint. The pretrained encoder produces a 512-dimensional CLS representation, while the gene-expression prediction (GEP) and masked value prediction for cell embedding (MVC) heads form the common objective structure used for continual learning and batch-correction training. Continual learning reconstructs masked gene expression through the GEP and MVC objectives. For batch correction, these prediction heads are additionally conditioned on batch-label information and combined with domain-specific batch normalization, elastic cell similarity, and domain-adversarial learning. The fused attention projection was separated into query, key, and value projections to connect the model to Hugging Face PEFT.

**Geneformer.** Geneformer was implemented using revision `04c2b2e8` of the official repository at `https://huggingface.co/ctheodoris/Geneformer` and the Geneformer V2-104M checkpoint. Following the V2 model configuration, cell representations were defined as the 768-dimensional CLS hidden states from the second-to-last Transformer layer. Continual learning used the pretrained BERT masked-language-modeling head to recover masked gene tokens, while sequence-control tokens were retained to preserve the input structure. Hugging Face PEFT methods were connected directly to the query and value projections of the Transformer attention layers.

**scFoundation.** The scFoundation implementation used revision `397631c4` of the official repository at `https://github.com/biomap-research/scFoundation` and its publicly released `cell` and `gene` checkpoints. The `cell` checkpoint was used for cell representation extraction, cell-type classification, and the custom MIL task, whereas perturbation prediction used the encoder–decoder structure provided by the `gene` checkpoint. Depending on the pooling setting, the model generated a 768-dimensional `max` representation or a 3,072-dimensional `all` representation. We added an AnnData-based Dataset and DataLoader to support batched execution while retaining the original cell-representation procedure. The RDA-MAE objective described in the original study and released configuration was implemented for continual learning. During mixed-precision profiling, the specified precision was applied to the encoder, whereas the large

Performer decoder was retained in FP32 for numerical stability. The fused attention projection was separated into query, key, and value projections for compatibility with Hugging Face PEFT.

**UCE.** UCE support was developed using revision `9c416007` of the official repository at `https://github.com/snap-stanford/UCE` and the released 4-layer checkpoint and protein-embedding resources. Genes were represented using the released 5,120-dimensional ESM-2 protein embeddings, which were loaded as a fixed lookup table and remained frozen throughout all inference and training experiments. The 4-layer UCE encoder transforms these protein-level inputs into a 1,280-dimensional CLS cell representation. Because the official implementation focuses on zero-shot representation extraction and does not provide a trainable classification architecture, we added a one-layer MLP that maps the cell representation directly to the target classes. The masked binary-expression prediction objective used for continual learning was implemented according to the original study. To support Hugging Face PEFT, the fused multi-head attention projection was converted into separate query, key, and value projections.

**Nicheformer.** Nicheformer was incorporated using revision `485cadbc` of the official repository at `https://github.com/theislab/nicheformer` and revision `0ba4ba16` of the Hugging Face model repository at `https://huggingface.co/theislab/Nicheformer`. The model represents each cell as a 512-dimensional vector obtained by averaging the valid gene-token hidden states. Following the released input procedure, gene expression was normalized using the technology-mean resource selected for each dataset. Assay context was specified either globally through the model configuration or, when corresponding annotations were available, for each cell through a designated AnnData `obs` column. Continual learning used the masked-token prediction head included in the released model. The fused attention projection was separated into query, key, and value projections to support Hugging Face PEFT.

#### Method S2. Model-specific pretraining objectives

For training-mode resource profiling, continual learning was performed using the self-supervised objective associated with each scFM. The model-specific input corruption, prediction module, and loss used in the current scUnify implementation are described below.

**scGPT gene-expression prediction and masked value prediction.** scGPT receives gene tokens together with binned expression values and replaces the expression values at selected positions with its mask value. Let $\mathcal{M}$ denote these masked positions, $x_i$ the original binned expression value at position $i$, and $\hat{x}_i^{\mathrm{GEP}}$ the output of the gene-expression prediction (GEP) decoder from the corresponding encoder hidden state. GEP minimizes the mean squared reconstruction error over the masked positions.

$$\mathcal{L}_{\mathrm{GEP}} = \frac{1}{|\mathcal{M}|} \sum_{i \in \mathcal{M}} \left(\hat{x}_i^{\mathrm{GEP}} - x_i\right)^2 . \tag{1}$$

The masked value prediction for cell embedding (MVC) decoder combines the cell representation with the corresponding gene-token embedding and predicts the same masked expression targets. Denoting its prediction by $\hat{x}_i^{\mathrm{MVC}}$, the MVC objective is

$$\mathcal{L}_{\mathrm{MVC}} = \frac{1}{|\mathcal{M}|} \sum_{i \in \mathcal{M}} \left(\hat{x}_i^{\mathrm{MVC}} - x_i\right)^2 . \tag{2}$$

The total objective used for continual learning was

$$\mathcal{L}_{\mathrm{scGPT}} = \mathcal{L}_{\mathrm{GEP}} + \mathcal{L}_{\mathrm{MVC}}. \tag{3}$$

**Geneformer masked language modeling.** Geneformer continual learning used the masked language modeling (MLM) head of the released BERT model. Gene tokens were selected dynamically from each input sequence, whereas special tokens such as <cls>, <eos>, and <pad> were excluded from masking. Selected gene tokens were either replaced with the mask token, replaced with a randomly sampled vocabulary token, or retained without modification. Let $\mathcal{M}$ denote the selected positions, $t_i$ the original gene token at position $i$, and $\widetilde{\mathbf{t}}$ the resulting MLM-corrupted input sequence. The objective was the mean cross-entropy over the selected positions.

$$\mathcal{L}_{\mathrm{Geneformer}} = -\frac{1}{|\mathcal{M}|} \sum_{i \in \mathcal{M}} \log p_\theta\Big(t_i \mid \widetilde{\mathbf{t}}\Big) . \tag{4}$$

**scFoundation RDA-MAE.** scFoundation continual learning used the read-depth-aware masked autoencoder (RDA-MAE) objective. For a cell with raw count vector $\mathbf{C}$, the encoder input was generated through stochastic read-depth down-sampling. For cells with at least 1,000 total counts, down-sampling was applied with probability 0.5 by drawing $b \sim \mathrm{Beta}(2, 2)$ and sampling $C_i^{\mathrm{in}} \sim \mathrm{Binomial}(C_i, b)$ for each gene. When down-sampling was not selected or the total count was below 1,000, the original counts were retained. The resulting input and original target were each normalized to 10,000 counts per cell and log1p-transformed. A subset of gene positions was then selected and corrupted to construct the masked input, which was passed through the scFoundation encoder and Performer decoder to reconstruct expression values for 19,264 genes. Let $\mathcal{M}$ denote the selected positions, $X_i$ the target expression, and $P_i$ the reconstructed expression at position $i$. The RDA-MAE loss was calculated as the mean squared reconstruction error over the selected positions.

$$\mathcal{L}_{\text{RDA-MAE}} = \frac{1}{|\mathcal{M}|} \sum_{i \in \mathcal{M}} (P_i - X_i)^2 \,. \tag{5}$$

**UCE masked binary-expression prediction** UCE continual learning used masked binary-expression prediction. For each cell, a subset of expressed genes was first removed, and the remaining expressed genes were used to construct a cell sentence from which the UCE encoder generated a cell representation. The removed expressed genes served as positive queries, whereas genes not expressed in the cell were sampled as negative queries. When the number of removed genes was smaller than the configured number of positive queries, all removed genes were included and the remaining positive queries were sampled from the expressed genes. For each query gene $j$, its fixed ESM-2 protein embedding was transformed by the released gene-embedding layer and concatenated with the cell representation. The UCE binary decoder then used this joint representation to produce a logit $a_j$ and the corresponding expression probability $p_j = \sigma(a_j)$. Let $\mathcal{Q}$ denote the combined set of positive and negative query genes and $y_j \in \{0, 1\}$ the observed expression label of gene $j$. The objective was calculated as binary cross-entropy over the query genes.

$$\mathcal{L}_{\text{UCE}} = -\frac{1}{|\mathcal{Q}|} \sum_{j \in \mathcal{Q}} \left[ y_j \log p_j + (1 - y_j) \log(1 - p_j) \right] . \tag{6}$$

**Nicheformer masked-token prediction.** Nicheformer continual learning used the masked-token prediction head included in the released Hugging Face model. scUnify supplied the unmasked sequence containing context and ranked gene tokens, after which the model's internal masking procedure selected eligible tokens and generated the corrupted sequence. Let $\mathcal{M}$ denote the selected positions, $t_i$ the original token, and $\widetilde{\mathbf{t}}$ the corrupted input sequence. The prediction head mapped the Transformer hidden state at each selected position to a distribution over the Nicheformer vocabulary, and the objective was

$$\mathcal{L}_{\text{Nicheformer}} = -\frac{1}{|\mathcal{M}|} \sum_{i \in \mathcal{M}} \log p_\theta\Big(t_i \mid \widetilde{\mathbf{t}}\Big) \,. \tag{7}$$

### Supplementary Figures

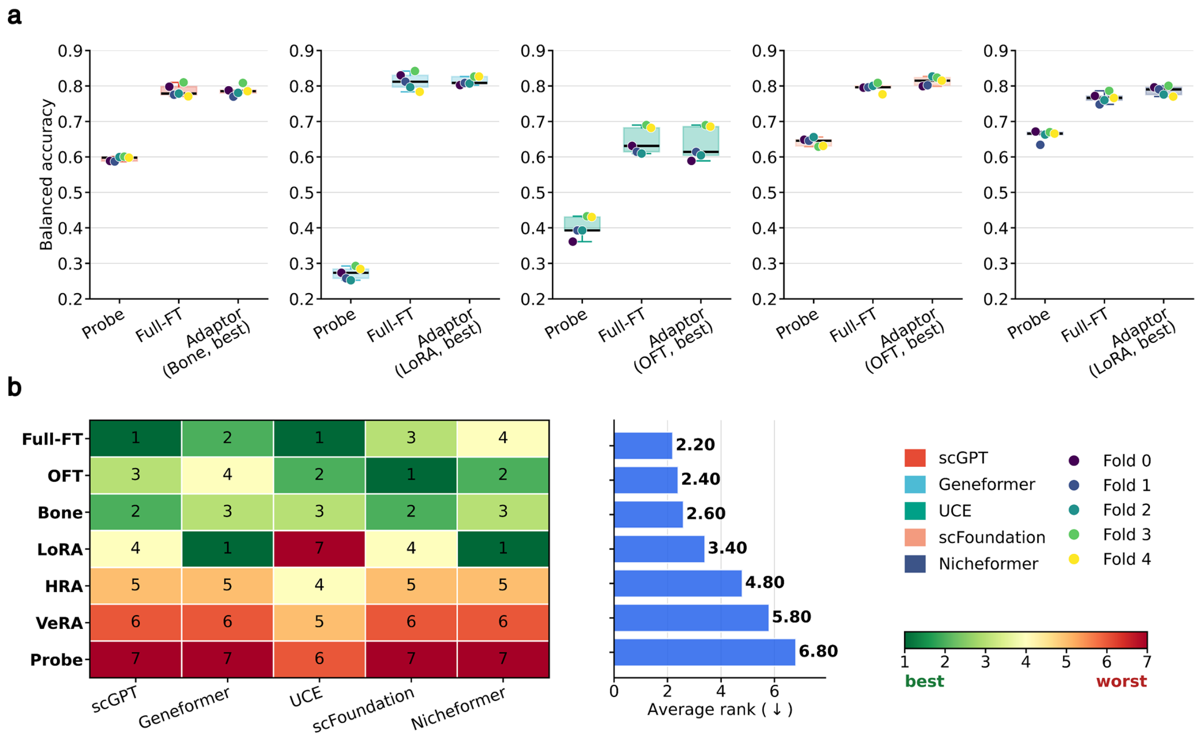

Fig. S1. **Standardized cell-type classification across five scFM backbones.** All five backbones were coupled to a common one-layer classifier and evaluated on the NSCLC dataset. Task-level training settings were held consistent across backbones within each adaptation condition. (a) Five-fold balanced accuracy for probing, Full-FT, and the PEFT method with the highest mean balanced accuracy for each backbone. Boxes summarize the distribution across folds, and colored points denote individual folds. (b) Within-backbone ranks of probing, Full-FT, LoRA, OFT, HRA, VeRA, and Bone (left) and their average ranks across the five backbones (right). Lower ranks indicate better performance.

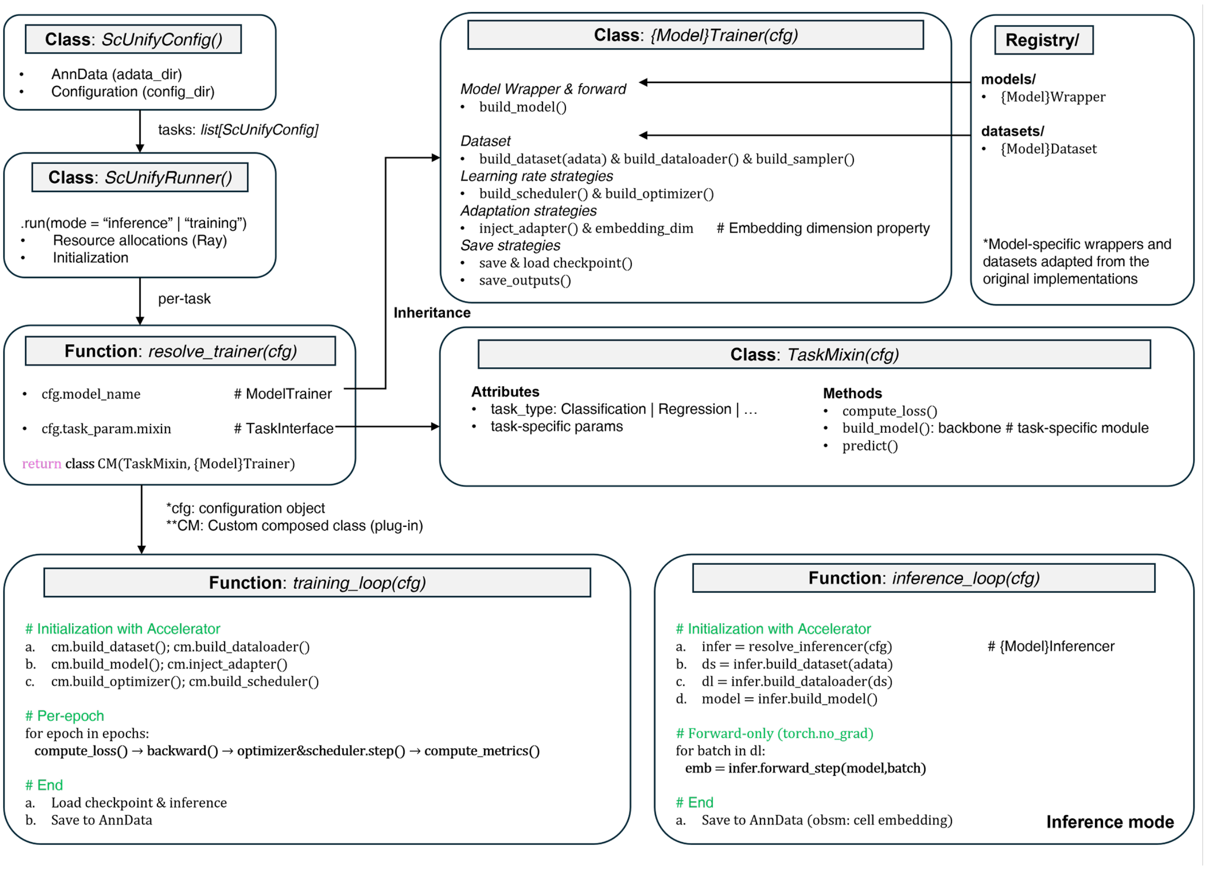

Fig. S2. **Architecture and execution flow of scUnify.** An scUnify run is defined by an AnnData file and a configuration interpreted by `ScUnifyConfig` and executed by `ScUnifyRunner`. For training, `resolve_trainer` dynamically composes the selected model-specific trainer and downstream-task mixin. The resulting trainer obtains model wrappers and datasets from the registry and provides the model, task objective, adaptation, optimization, output, and checkpoint procedures used by the training loop. Inference follows a separate forward-only path that loads the selected model-specific inferencer and stores the resulting cell embeddings in AnnData.

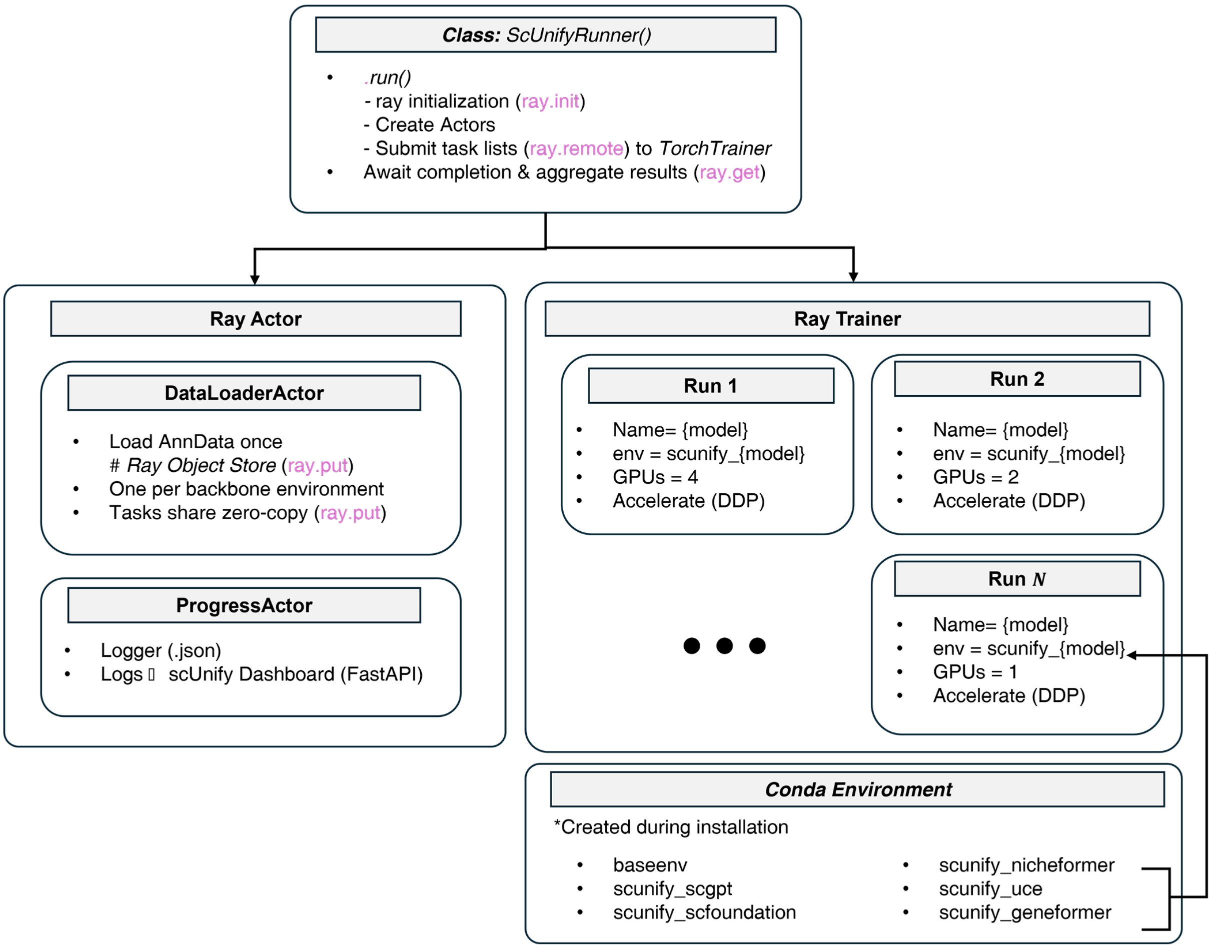

Fig. S3. **Ray-based organization of the scUnify core system.** `ScUnifyRunner` initializes Ray, creates the data and progress actors, submits individual runs to Ray TorchTrainer, and aggregates their outputs. `DataLoaderActor` loads AnnData into the Ray Object Store so that runs using the same dataset can share a common reference, while the progress actors record run status and progress for the dashboard. Each run is assigned its requested CPU and GPU resources, executed in the model-specific Conda environment, and uses Hugging Face Accelerate for Distributed Data Parallel execution when multiple GPUs are assigned.

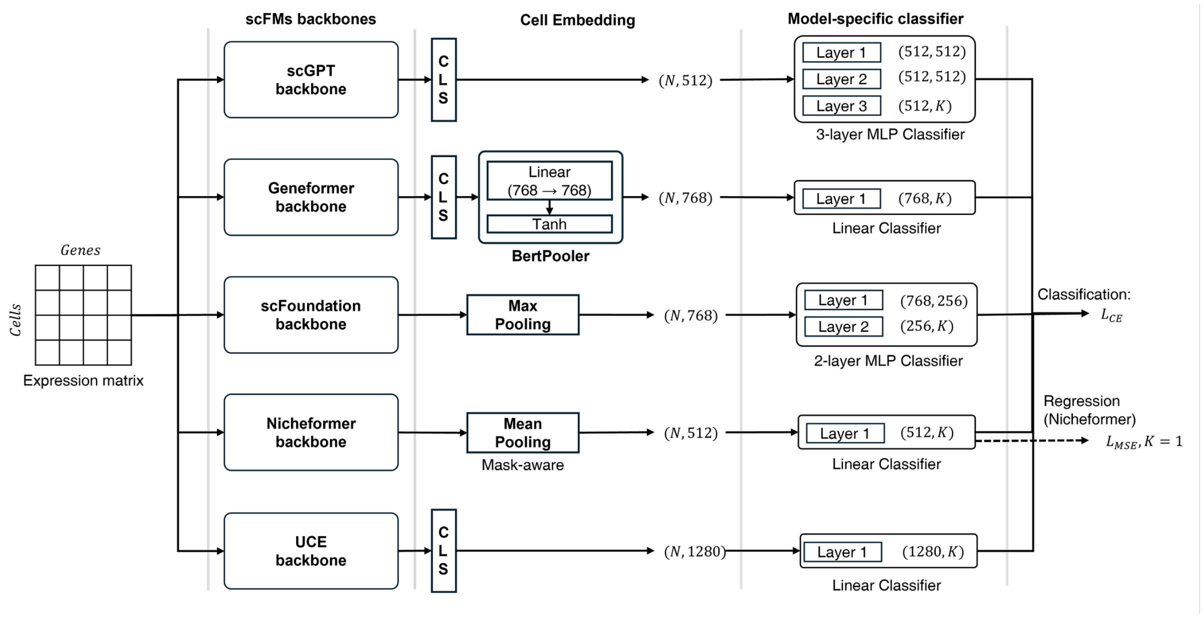

Fig. S4. **Model-specific architectures used for the built-in cell-type classification and Nicheformer regression tasks.** Geneformer, scGPT, scFoundation, Nicheformer, and UCE generate cell representations using their respective CLS extraction or pooling procedures. Each representation is passed to the downstream prediction architecture derived from the corresponding original implementation where available; for UCE scUnify uses the added one-layer classifier. The Nicheformer linear prediction head is used for both classification and niche-density regression, with a scalar output for regression.

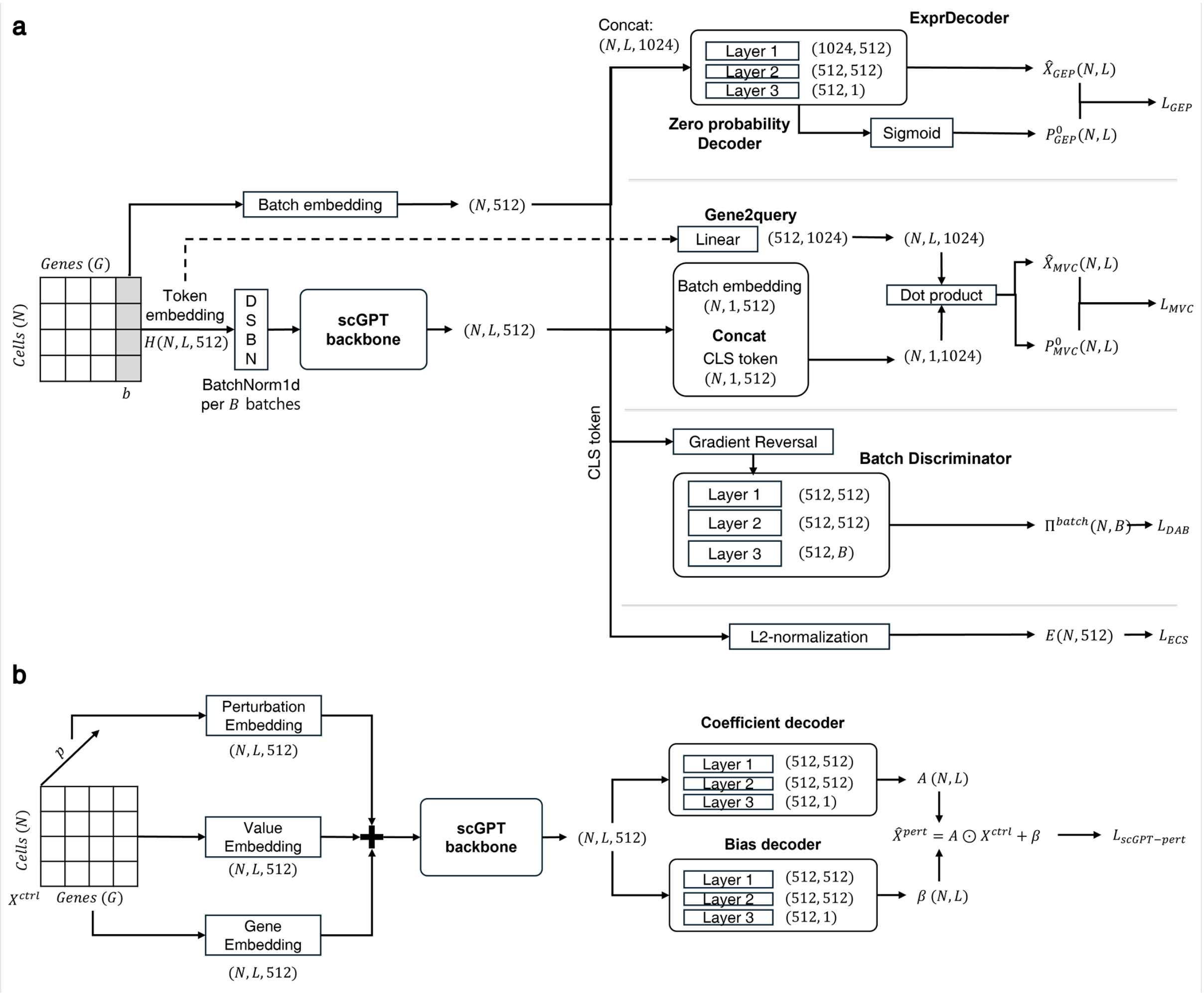

Fig. S5. **Architectures of the scGPT built-in downstream tasks.** (a) For batch correction, domain-specific batch normalization (DSBN) conditions the input on its batch label before processing by the scGPT backbone. Encoder hidden states and the batch embedding are used by the gene-expression prediction and zero-probability decoders, while the CLS representation supports masked value prediction from the cell representation, domain-adversarial batch classification through gradient reversal, and elastic cell-similarity regularization after L2 normalization. (b) For perturbation prediction, gene, control-expression value, and perturbation-condition embeddings are combined and processed by the scGPT backbone. The resulting hidden representations are passed to separate coefficient and bias decoders, whose outputs are combined with the control expression to predict the perturbed gene-expression profile.

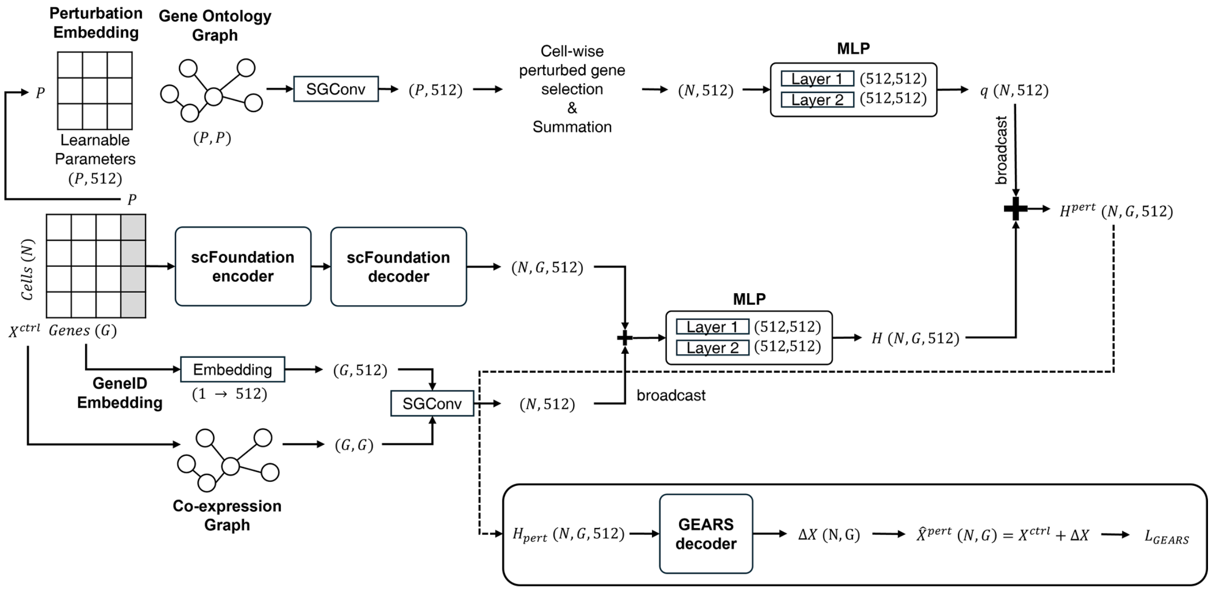

Fig. S6. **Architecture of scFoundation perturbation prediction.** Control expression is processed by the scFoundation encoder and decoder to generate gene-level representations, which are propagated through the gene co-expression graph using simplified graph convolution (SGConv). Learnable perturbation-gene embeddings are propagated through the gene-ontology graph and aggregated for each perturbation condition. The gene and perturbation representations are then combined by the GEARS decoder to predict perturbation-induced expression changes.
